# SODA: Multi-locus species delimitation using quartet frequencies

**DOI:** 10.1101/869396

**Authors:** Maryam Rabiee, Siavash Mirarab

## Abstract

**Motivation:** Species delimitation, the process of deciding how to group a set of organisms into units called species, is one of the most challenging problems in evolutionary computational biology. While many methods exist for species delimitation, most based on the coalescent theory, few are scalable to very large datasets and methods that scale tend to be not accurate. Species delimitation is closely related to species tree inference from discordant gene trees, a problem that has enjoyed rapid advances in recent years.

**Results:** In this paper, we build on the accuracy and scalability of recent quartet-based methods for species tree estimation and propose a new method called SODA for species delimitation. SODA relies heavily on a recently developed method for testing zero branch length in species trees. In extensive simulations, we show that SODA can easily scale to very large datasets while maintaining high accuracy.

**Availability:** The code and data presented here are available on https://github.com/maryamrabiee/SODA

## 1 Introduction

Evolution results in diversity across species and diversity within the same species, in ways that can make it difficult to distinguish species. Definitions of what constitutes a species are varied Coyne and Orr, 2004 and subject to debate. Nevertheless, many biological analyses depend on our ability to define and detect species. Assigning groups of organisms into units called species, a process called species delimitation, is thus necessary but remains challenging (Carstens *et al*., 2013). Among varied species concepts, the most commonly used for Eukaryotes is the notion that individuals within a species should be able to mate and reproduce viable off-springs.

A wide range of species delimitation methods exist. More traditional methods simply relied on the mean divergence between sequences (e.g., Hebert *et al*., 2004; Puillandre *et al*., 2012) or patterns of phylogenetic branch length (Zhang *et al*., 2013; Fujisawa and Barraclough, 2013; Esselstyn *et al*., 2012) in marker genes or concatenation of several markers (e.g. Pons *et al*., 2006). Due to limitations of marker genes (Hudson and Coyne, 2002), many approaches to species delimitation have moved to using multi-locus data that allow modeling coalescence within and across species (Knowles and Carstens, 2007; Yang and Rannala, 2010; O’Meara, 2010), not to mention more complex processes such as gene flow (e.g., Leaché *et al*., 2019). Modeling coalescence allows methods to account for the fact that across the genome, different loci can have different evolutionary histories, both in topology and branch length (Maddison, 1997). Species delimitation is often studied using the Multi-species Coalescent (MSC) model (Pamilo and Nei, 1988; Rannala and Yang, 2003). In this model, individuals of the same species have no structure within the species and thus their alleles coalesce completely at random. Coalescence is allowed to happen deeper than the first opportunity, producing gene tree discordance due to Incomplete Lineage Sorting (ILS). In this context, given a set of sampled individuals, delimitation essentially requires inferring gene trees, one per locus, and detecting which delimitation is most consistent with patterns of coalescence observed in the gene trees.

Many methods for species delimitation under the MSC model exist, but they tend to suffer from one of two limitations. The most accurate set of methods are the Bayesian MCMC methods that infer gene trees, (optionally) species trees, and species boundaries (e.g., BPP (Yang and Rannala, 2010, 2014a), ABC (Camargo *et al*., 2012), and STACEY (Jones, 2017)). Other Bayesian methods use biallelic sites (Leaché *et al*., 2014), incorporate morphological data (Solís-Lemus *et al*., 2015), or use structure (Huelsenbeck *et al*., 2011). These methods, however, are typically slow and cannot handle even moderate numbers of samples (Fujisawa and Barraclough, 2013; Xu and Yang, 2016). For example, Musher and Cracraft (2018) had to divide their dataset of 62 individuals into six subsets, and Oliveira *et al*. (2015) used a subsample of 20 out of 137 to use BPP to avoid mixing problems that usually happen with large datasets; running BPP on a dataset of 40 populations in our study needed 36 hours of running time. A second class of methods (e.g., SpedeSTEM (Ence and Carstens, 2011)) rely on a three-step approach: first infer gene trees, then, date gene trees so that they all become ultrametric (i.e., have a unique root to tip distance), and finally, use ML calculation of alternative delimitations under the MSC model to decide species boundaries. These methods have been less accurate than Bayesian methods and their reliance on ultrametric trees make them hard to use for datasets where rates of evolution change substantially across the tree (Camargo *et al*., 2012). Yet other methods (e.g., O’Meara, 2010; Zhang and Cui, 2010) rely only on input gene tree topologies, as we do.

In this paper, we introduce a new species delimitation approach called SODA that builds on the success of our species tree inference tool ASTRAL (Mirarab *et al*., 2014a; Mirarab and Warnow, 2015; Zhang *et al*., 2018). As a statistically consistent method, ASTRAL infers a species tree from a collection of gene tree topologies (ignoring branch lengths) based on the principle that the most frequent unrooted topology for each quartet of species is expected to match the species tree (Allman *et al*., 2011). Thanks to its accuracy and scalability, ASTRAL has been widely adopted for species tree inference. In a recent paper that extended ASTRAL to multi-individual data, we observed that if species boundaries are ignored, ASTRAL most often recovers individuals of the same species as monophyletic (Rabiee *et al*., 2019). This result suggests a species delimitation method: Infer an ASTRAL tree with all individuals and use patterns of quartet trees mapped onto that species tree to decide where coalescence is completely random and where it is not; these boundaries readily define species. By relying on quartet frequencies and ASTRAL machinery, SODA is able to handle very large datasets with short running times. We describe SODA in detail and then perform two sets of extensive simulation studies to evaluate its accuracy and scalability.

## 2 Methods

### 2.1 Coalescent-based topology-based delimitation

We take a two-step approach to species delimitation and assume unrooted gene tree *topologies* are already inferred from sequence data. Thus, we are given a set of unrooted gene trees 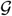 on individuals 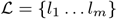. Ignoring errors in estimated gene trees, we assume these gene trees follow the MSC model. Optionally, we are given a partition of 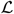 into populations 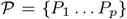; when 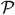 is not given, we define each individual as a singleton population. A partition of 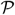 into 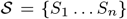 produces a mapping *r* : 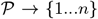 and by extension *q* : 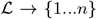 where *r*(*x*) and *q*(*y*) give a species index for 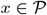 and 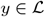.

We say a partition is coalescent-consistent if for any set 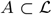 with at most one individual from each population and all individuals mapped to the same species 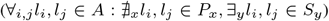, the distribution of 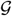 restricted to *S* is consistent with the neutral Kingman (1982) coalescence process. Because Kingman’s process is robust to subsampling, if a partition is coalescent-consistent, further breaking each species into smaller species would remain coalescent-consistent. Having two species that could be combined without violating the coalescent model is not justified under the MSC because the model provides no support for the division. Thus, we formulate coalescent-based species delimitation as the problem of finding a coalescent-consistent partition that is not a refinement of any other coalescent-consistent partitions. Our method seeks to solve this problem within further restrictions stated below.

This formulation follows the MSC model (free coalescent of lineages within species but constrained coalescence across the species), and thus shares its assumptions. The only population structure within the species that is modeled is the given structure 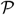, which is known *apriori*. Thus, it assumes that individuals selected from different populations of the same species evolve according to the neutral Wright-Fisher model, resulting in a distribution of gene trees within a species that follows the Kingman (1982) coalescent process. This assumption is most defensible when the populations within a species do not further have *strong* differentiation. MSC also assumes lineages sampled from different species do not coalesce more recently than their separation event (Fig. S8); thus, it ignores gene flow across species. While these assumptions can all be violated on real data, they provide a useful model that allow fast delimitation. We revisit these assumptions in the discussion session.

The branch lengths of gene trees can be modeled as a function of two processes: coalescent of lineages and changes in the mutation rate (Rannala and Yang, 2003). Dealing with these two processes simultaneously is a difficult computational challenge, motivating methods such as SpedeSTEM to take ultrametric (e.g., dated) gene trees as input and forcing Bayesian methods such as BPP to assume parametric rate models. In our work, we are after a fast delimitation method that can be applied to inferred non-ultrametric gene trees directly. To avoid complications of rate variations across lineages, we limit ourselves to gene tree topologies. To do so, we rely on the distribution of gene tree topologies under the MSC model, in particular for quartets of speices (Degnan and Salter, 2005).

Using gene tree topologies, however, has a limitation. Examining the distribution of tree topologies requires at least three lineages. Thus, two species each with an individual sampled and one species with two individuals cannot be distinguished by topology alone. This forces us to assume that in the correct delimitation, each species has more than one individual sampled (i.e., ∀_i_ |*S_i_*| ≥ 2).

### 2.2 SODA Algorithm

A central concept in MSC is the “extended species tree”, as defined by Allman *et al*. (2011). Let *T*\* be the true species tree on the leafset 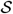. The extended species tree 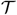 is a rooted tree labeled by 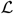, built by adding to each leaf of *T*\* all individuals corresponding to that species as a polytomy (Fig. 1); i.e., for leaf 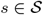 of *T*, add a child for every 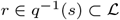. When populations are known *apriori*, we can similarly define the extended species tree 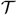 as a rooted tree labeled by 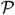, built by adding to each leaf of *T*\* all populations corresponding to that species as a polytomy.

**Fig. 1.**
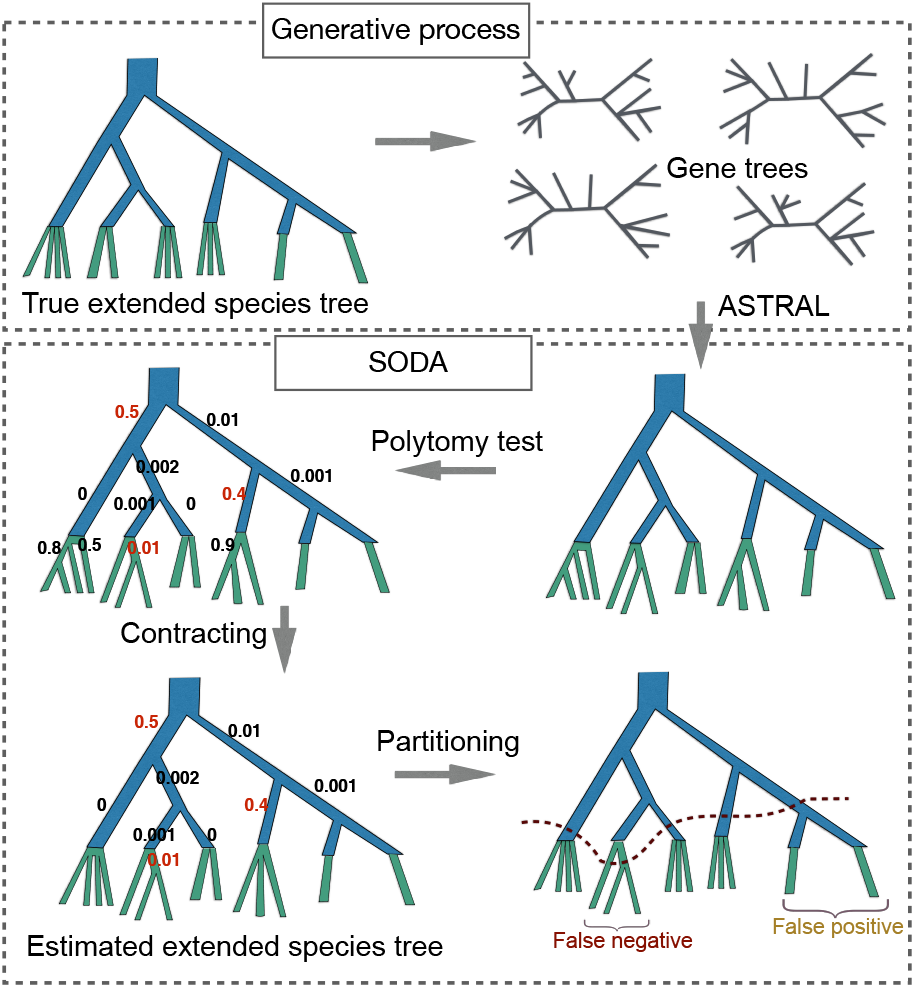
True extended species tree generates gene trees under the MSC model. From unrooted gene tree topologies (inferred from sequence data), SODA first estimates a guide tree using ASTRAL and then test the null hypothesis that each branch has length zero length, obtaining a p-value (middle left). *p*-values may result in FP or FN rejection or retention of the null (red *p*-values). SODA then contracts branches where the null is retained as long as contracting them does not contradict with the monophyly of species defined by branches where the null hypothesis is rejected; thus, we keep some branches with high *p*-value (e.g., those with *p*-value 0.4 and 0.5) in a way that ensures the resulting tree can be an extended species tree (bottom left). The inferred extended species tree can be cut at branches above the terminal branches to define species. The result could include both false positives and false negative delimitations. However, we note that some errors in *p*-values (0.5 and 0.4) do not result in error in delimitation.

**Algorithm 1.**
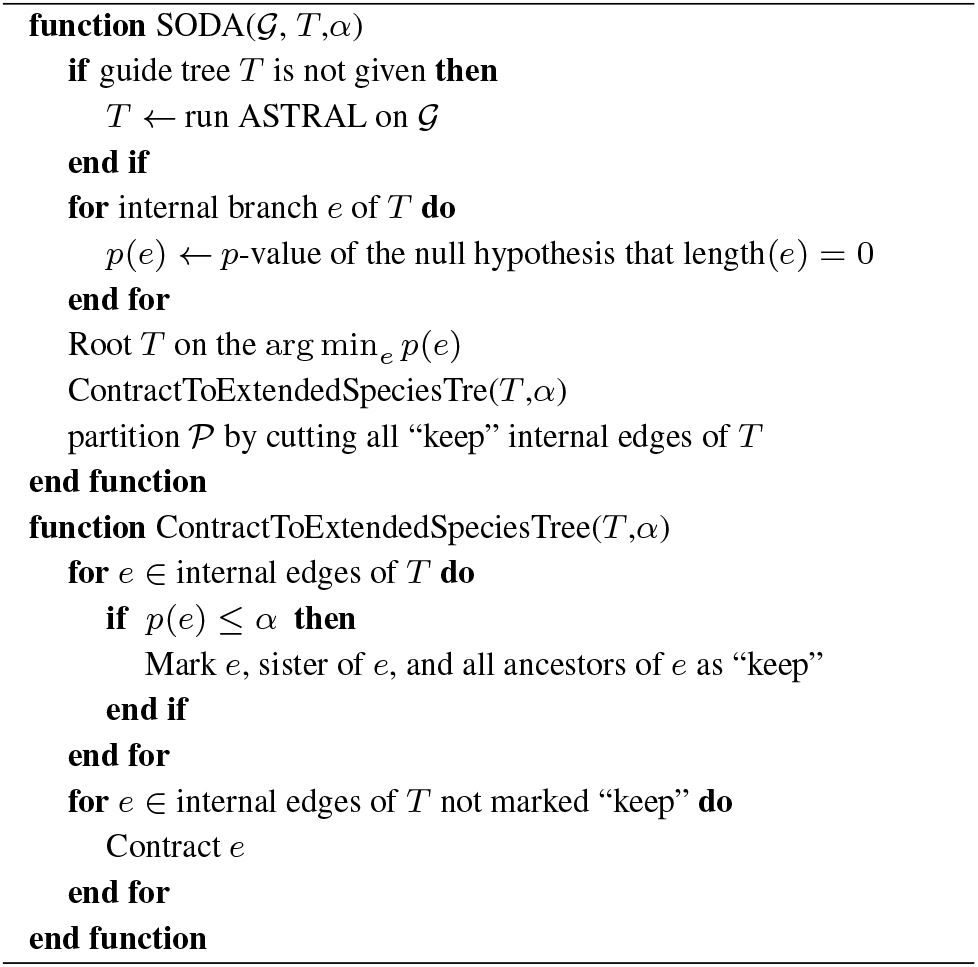
SODA Algorithm

Our species delimitation method, which we name Species bOundry Delimitation using Astral (SODA), is shown in Algorithm 1. Its inputs are a set of gene tree topologies and a significance level *α*, described below. SODA first infers (or takes as input) a guide tree *T*. The guide tree (a concept introduced by Yang and Rannala (2010)) is a phylogenetic tree with leaves set to known populations (i.e., the most divided possible delimitation). It assumes *T* is a *resolution* of the true extended species tree 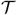, meaning that it includes all branches of the species tree and has arbitrarily relationships between populations of the same species. SODA then contracts *some of the* branches of *T* as long as it cannot reject the null hypothesis that they have length zero under the MSC model using *α* as the confidence level for statistical tests; the contracted tree is an estimated extended species tree 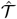. Finally, SODA cuts internal branches of the 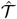 to cluster species (Fig. 1). Below, we describe each step in more details.

#### Guide Tree

SODA needs a (potentially unrooted) guide tree *T* on the leafset 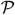. If not provided by the user, we infer the guide tree by running ASTRAL-III on 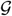; note that using the multi-individual version of ASTRAL (Rabiee *et al*., 2019), we can ensure that the tree generated by ASTRAL is labelled by 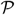, as opposed to original individuals (if different). We assume that *T* is a resolution of the extended species tree; thus, the accuracy of this guide tree is important.

#### Polytomy Test

For each branch of *T*, we next test the null hypothesis that it has zero length in coalescent units. If we cannot reject this null hypothesis, the branch can be collapsed to obtain a polytomy, helping us to obtain 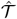. We use a recent test proposed by Sayyari and Mirarab (2018) that relies on a classic result: Under the MSC model, across gene trees, the frequencies of the three resolutions for each quartet around a given branch in the species tree are equal if and only if that branch has length zero (Pamilo and Nei, 1988; Allman *et al*., 2011). Moreover, gene trees are assumed independent. Thus, under the null hypothesis of a polytomy, the frequency of quartet topologies around each branch should follow a multinomial distribution with three categories each with probability 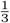. Whether observed frequencies of the three possible topologies for a quartet follow an equiprobable multinomial distribution can be tested using a Chi-Squared test, which is what the method of Sayyari and Mirarab (2018) uses. To achieve scalability, this method treats branches independently (based on the locality assumption of Sayyari and Mirarab (2016)) and takes the average of quartet frequencies for all quartets around each branch (which can be done easily in *O*(*n*^2^*k*) time). The test assumes the input the gene tree set is an error-free random sample generated by the MSC model from the true species tree. It produces one *p*-value per *internal* branch.

#### Rooting *T*

We need to root *T* (if not rooted) such that each species becomes monophyletic. We simply root *T* at the edge with the minimum *p*-value. Note that our goal is not to find the correct root because we do not need the correct rooting in the next steps. We only need the tree to be rooted on *any* internal branch of the extended species tree. The highest statistical confidence for having a positive length is achieved by the branch with the lowest *p*-value; thus we can root here.

#### Infer extended species tree

To obtain 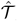, we contract some of the branches of *T* where a zero length null hypothesis cannot be rejected at a user-specificed level *α*. When the null hypothesis is rejected for a branch *e*, we marked *e* as being part of 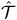 (i.e., *keep* in Alg. 1). Parameter *α* can be adjusted for controlling how aggressively SODA divides species. Increasing *α* results in rejecting more null hypotheses and hence, dividing individuals into more species. To ensure that we can get a valid extended species tree (with monophyletic species), we need polytomies to form only above terminal branches. Thus, in addition, we mark the sister edge of *e* and all its ancestor edges as belonging to 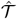.

#### Partitioning

Given 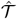, the partition is obtained by cutting remaining internal branches of 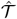. The partition produced would be identical if we only cut internal branches that have at least one terminal branch as child.

The accuracy of the algorithm, in addition to assumptions made by our problem formulation, depends on the accuracy of the statistical test. We formalize this notion in three claims (proofs in supplementary material).

##### Claim 1.

*Assuming* (*i*) *gene trees* 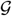 *are generated under the MSC model on an extended species tree* 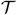, (*ii*) *the guide tree T* (*e.g., ASTRAL tree*) *is a resolution of* 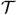, *and* (*iii*) *the hypothesis testing has no false positive* (*FP*) *or false negative* (*FN*) *errors, the SODA algorithm returns the correct extended species tree* 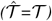. *Additionally, it will correctly delimitate species if all species are sampled more than once*.

This claim provides a reassuring positive result, but only under strong assumptions, most notably, that the test is perfect. However, errors in the test *can* lead to errors.

##### Claim 2.

*Given a guide tree T thatresolves the tree* 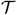, *SODA incorrectly divides a species S into multiple species* (*i.e., a false negative error*) *if and only if the zero length hypothesis testing results in an FP error for one of the branches under the clade defined by S on T*.

Thus, FPs in the hypothesis test always result in division of a species; however, an FP does not always divide *S* into exactly two parts. For example, if *T* has a caterpillar (a.k.a ladder-like) topology on *S*, an FP on the branch above the cherry (i.e., a node with two leaf children), leads to each remaining individual being marked as a species.

##### Claim 3.

*Given a guide tree T that resolves* 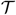, *SODA incorrectly combines individuals from two species S*_1_ *and S*_2_ *into one species* (*a false positive error*) *under one of these two conditions. 1) S*_1_ *and S*_2_ *each have one sampled individuals and form a cherry. 2) The hypothesis testing has an FN error for all branches of T below the LCA of S*_1_ *and S*_2_. *This condition requires FN errors for two or more branches if neither species is a singleton*.

To combine two species incorrectly, we must fail to correctly mark all branches below their LCA; else, one of the branches below the LCA would be cut, which would prevent the FP. Thus, our approach is tolerant of some FN errors in the hypothesis test (e.g., *p*-value 0.4 in Fig. 1).

## 3 Experimental setup

### 3.1 Datasets

We used two simulated datasets, both generated using Simphy (Mallo *et al*., 2016). One dataset is large and allows us to evaluate SODA on datasets where other methods cannot run whereas the other dataset is small and enables us to compare our method to slower Bayesian methods.

#### Large dataset

We reuse a 201-species “homogeneous” dataset that we have previously simulated (Rabiee *et al*., 2019) using SimPhy to generate species trees based on the birth/death model and gene trees under the MSC model. Each species includes 5 individuals except the singleton outgroup (1001 in total). We have three model conditions (50 replicates each) with medium, high, or very high levels of ILS, with maximum tree height set to 2M, 1M, and 0.5M generations, respectively. The mean quartet score of true gene trees versus the species tree (i.e., the proportion of quartet trees that are shared between the two trees), are 0.78, 0.62 and 0.50 for these model conditions (Fig. S5). Note that a quartet score of 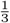 indicate random trees, and even a value as low as 0.5 indicates very high levels of ILS. The proportions of branches in the true extended species tree missing from gene trees are 0.40, 0.57 and 0.77. Each replicate has 1000 genes, which we down-sample randomly to 500, 200, and 100. To evolve nucleotide sequences down the gene trees, we use INDELible (Fletcher and Yang, 2009) with the GTR+Γ model of sequence evolution with randomly sampled sequence lengths from a LogNormal distribution (empirical mean=721). The gene trees are estimated using FastTree based on the alignments (Price *et al*., 2010) and estimated gene trees are used throughout the experiments in this paper.

The average gene tree RF error for the three model condition are 0.25, 0.31 and 0.42 with large variance (Fig. S5).

#### Small dataset

We simulate a new dataset using SimPhy with 20 replicates, each only 4 species, 10 individuals per species, and 1000 genes per replicate (commands shown in Appendix 1). The tree height is set to 200,000 generations, the population size is drawn uniformly between 10,000 and 500,000, and species trees are generated using birth-only model with rate=0.00001. These settings lead to high level of ILS, capturing a scenario close to where species delimitation is of relevance; the quartet score of true gene trees versus the true species tree is 0.76 and 65% of extended species tree branches are on average missing from true gene trees. We deviate from ultrametricity by drawing rate multipliers for species and genes from Gamma distributions, with LogNormal priors on parameters of Gamma (Table S1). We simulate 1000bp alignments on each gene tree using INDELible (Fletcher and Yang, 2009) and estimate gene trees using FastTree based on the alignments (Price *et al*., 2010). The average gene tree error (normalized (Robinson and Foulds, 1981) distance (RF) between true and estimated gene trees) is 43%. Gene tree distance between pairs of individuals of each species had a wide range in our simulations, falling between 10^−5^ and 10^−2^ mutations per site in most cases (Fig. S2).

#### Empirical dataset

We study three biological datasets. To show applicability on large data, we use the dataset of Protea L. with recent radiations (Mitchell *et al*., 2017), which has sampled multiple individuals from 59 species of Protea and 6 outgroup species (a total of 163 tips) and obtained 498 low-copy, orthologous nuclear loci. Due to its size, this dataset is only analyzed using SODA. To be able to compare to other methods, we use a smaller dataset of lizards of the Australian wet tropics (AWT). Singhal *et al*. (2018) analyzed genetic data for individuals from three species groups – the *Carlia rubrigularis*, *Lampropholis coggeri*, and *Lampropholis robertsi* groups that split into 13 putative lineages. In total, there are 25 individuals, and 3320 loci across all individuals are sequenced using an exome capture approach. Gene trees and the species tree are estimated using STARBEAST2 v0.13.5 (Ogilvie *et al*., 2017), and delimitation results using STACEY (Jones, 2017) and BPP are available from the original study. We also study the human dataset analyzed by Jackson *et al*. (2017) comprising sequences from 50 loci (415 to 960 bp long) for four widely sampled groups of humans defined geographically (as original study states: these four groups are ethnically diverse and are not “populations” in any biological sense). The dataset includes 10 samples from each of Africa, Europe, and Asia, and 12 samples from South and Central America. Analyzing the human dataset, where we clearly know all individuals belong to one species, enables us to test the propensity of the method to lead to false positive delimitation.

### 3.2 Measures of accuracy

We evaluate accuracy using two measures.

**ROC.** Each pair of individuals is categorized depending on whether they are correctly grouped together (TP), correctly not grouped together (TN), incorrectly grouped together (FP), or incorrectly not grouped together (FN). We then show Recall = TP/(TP+FN) and FPR = FP/(TN+FP) on an Receiver Operating Characteristic (ROC) curve as we change *α* of SODA.
**Adjusted Rand Index** is a similarity measure between two partitions of a set also based on pairwise comparisons. We report ARI between the true species partition and the partition estimated by each method. The Rand (1971) index is 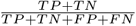. The adjusted rand index (ARI) adjusts RI for expected similarity of pairs according to a generalized hyper-geometric distribution that controls for the number of objects and classes (Hubert and Arabie, 1985). ARI equals 1 only for the correct partition and is close to 0 for a random partition.

### 3.3 Methods compared

#### BPP

We compare SODA to the widely used method Bayesian Phylogenetics and Phylogeography (BPP). BPP uses MCMC for inferring species boundaries directly from sequence alignments by sampling gene trees and other model parameters (e.g., rates) under the MSC model. We use BPP 4.1.4 and took advantage of its multi-thread version to be able to run it with up to 1000 genes. Provided with all 40 individuals of the small dataset that we sampled as 40 separate populations, BPP could not run to completion in 36 hours, perhaps because the set of possible delimitations was too large. To be able to test BPP, we use two subsampled sets. First, we randomly sample 4 individuals per species, and designate each as its own population, for a total of 16 populations. In the second scenario, from each species, we sample 7 individuals and randomly assigned them to 3 populations of sizes 2, 2 and, 3, comprising 12 populations in total. We use a uniform prior across all possible partitions on the resulting sets of populations. BPP calculates the posterior probability for each partition, and we use the delimitation with the highest posterior probability to measure the accuracy of the method.

We explore settings of BPP as follows. The total number of MCMC iterations is set to 208000 with the first 8000 discarded as burnin. We also run BPP with twice the number of iterations in one experiment with 500 genes to ensure convergence is not an issue. For the required species tree (similar to guide tree for SODA), we run BPP in two ways. By default, we provide BPP with the ASTRAL species tree (just as we do for SODA). The guide trees are rooted to match the true species tree. However, BPP is able to jointly infer species trees and species delimitation based on sequence alignments; we also run BPP with this co-estimation setting. This setting makes BPP 2 × slower (even though we only have four species), making it impractical for an extensive study. The priors for the inverse gamma distribution parameters, (*α, β*), were chosen to be (1.525,0.0001), (1.525,0.001) and (1.525,0.01) for population size *θ_s_* and twice the *β* for *τ_s_*. The mean of the distribution is set based on the true average of species pairwise distances in the gene trees. An example of a control file used for running BPP is given in Figure S1 for the full set of parameters.

#### SpedeSTEM

Given a set of rooted ultrametric gene trees, SpedeSTEM uses the STEM (Kubatko *et al*., 2009) algorithm to calculate the maximum likelihood species tree considering possible species tree and delimitation combinations and uses AIC to select among the models. SpedeSTEMv0.9 software includes a pipeline for inferring ultrametric rooted gene trees using paup* (Swofford, 2001). We used SpedeSTEMv0.9 to infer the rooted ultrametric input gene trees, which we then fed to the SpedeSTEMv2 software as input. We ran delimitation using theta values of 0.1 and 0.01, and sampling ratio of 1 (no subsampling). SpedeSTEMv2 also requires a putative assignment of populations to species. For this, we tried to assigning all populations to the same putative species or assigning all 12 populations to individual species. Results were similar and we report the latter strategy.

#### SODA

We implemented SODA in python using Dendropy (Sukumaran and Holder, 2010). We vary *α* between 0.005 and 0.5 but designate *α* = 0.05 as default. We infer the guide tree using ASTRAL-III on all 1000 genes. To study the impact of the guide tree, we also create the true extended species tree and resolve its polytomies randomly and use this guide tree as input to SODA. For tests with known populations, we inferred the guide tree using ASTRAL-multi, mapping leaves of the same population to a “species”, assigning each population to a separate species.

## 4 Results

### 4.1 Large simulated dataset

On the large dataset with 1001 individuals, SODA takes no more than 35 minutes (Table S2) and is highly accurate (Fig. 2). Given 1000 genes, default SODA (*α* = 0.05) is able to recover, on average, 183, 186, and 189 out of the 201 species entirely correctly for the three model conditions in decreasing order of ILS (Fig. 2a). The total number of species estimated by SODA ranges between 183 and 220 across all replicates with mean=208, which slightly over-estimates the correct number. The number of detected species increases as *α* increases; however, the number of correct species does not always increase. Reducing the number of genes reduces the number of correctly estimated species (down to 158 with 100 genes, very high ILS); however, it does not change the total number of species dramatically (for default *α*).

**Fig. 2.**
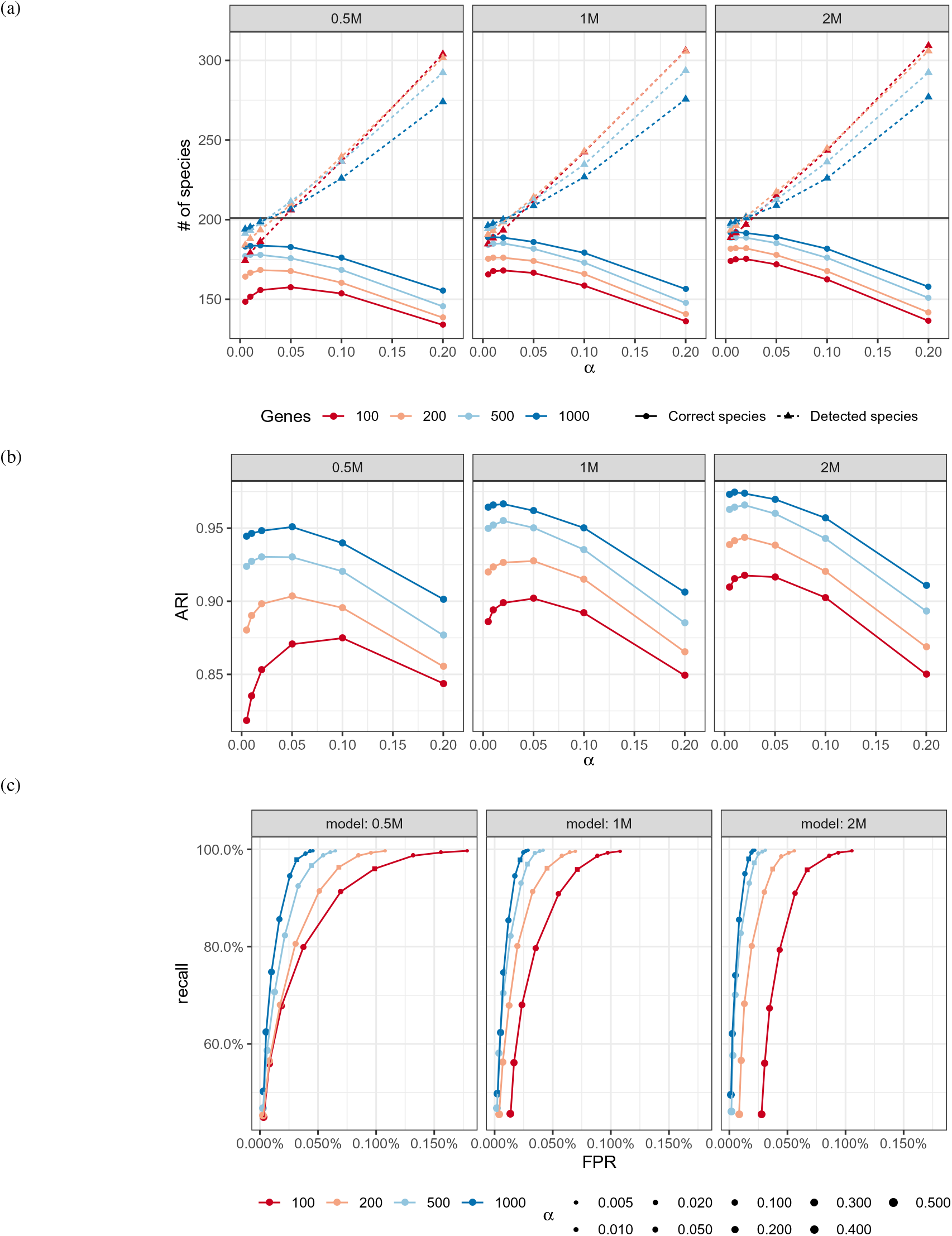
Accuracy of SODA on the large dataset. (a) We show the number of species that are completely correctly delimited (solid lines) and the total number of species found by SODA (dashed lines). Results divided into three model conditions with very high ILS (0.5M), high ILS (1M) and moderate ILS (2M) as we change *α* (x-axis) and the number of genes (colors). We clip *α* at0.2 but show full results in Figure S3. (b) ARI (y-axis) shows the accuracy of SODA. (c) ROC showing recall versus False Positive Rate (FPR) for all model conditions and different choices of *α* (dot size). The default value shown as a square.

A substantial portion of mistakes made by SODA are related to the guide tree. The ASTRAL tree failed to recover on average 7, 5 and 4 species as a monophyletic clade in these three model conditions, and these species could never be recovered correctly by SODA. On average, the estimated guide tree by ASTRAL (on 500 genes) missed 8%, 6%, and 4% of branches in the true extended species tree for high, moderate, and low levels of ILS. The *estimated* extended species tree that SODA outputs has an RF distance of 5%, 4%, and 2% to the true extended species tree.

Examining pairs of individuals, we observe very high accuracy. With *α* = 0.05, the ARI ranged between 0.95 and 0.97 for our three conditions given 1000 genes and between 0.87 and 0.92 when given as few as 100 genes (Fig. 2b). Reducing *α* to 0.005 or increasing it to 0.1 can reduce or increase ARI slightly; however, increasing *α* beyond 0.1 can quickly lead to substantial reductions in ARI (Fig. 2b). The best choice of *α* is always between 0.01 and 0.1, but 0.05 is never far from optimal, motivating us to use it as default. Since our simulated replicates are very heterogeneous in terms of gene tree estimation error, we can also examine the impact of mean gene tree error on the accuracy of SODA. Except for an outlier replicate with very high gene tree error and low ARI < 0.9, we do not detect a strong correlation between gene tree error and accuracy (Fig. S6). However, the guide tree and the extended species tree *are* impacted by increased gene tree error (Fig. S7). As the increased error does not impact delimitation, the increased error of the guide tree must be concentrated on deep branches that do not impact delimitation.

The trade-off between precision and recall with different choices of *α* can be examined using the ROC curve (Fig. 2c). With *α* ≤ 0.05, recall is always 96% orhigher, andis often close to 100% with *α* ≤ 0.01.The FPR, however, is strongly impacted by the number of genes. For example, with default *α*, FPR is never more than 0.03% with 1000 genes but increases to 0.98% with 100 genes. Reducing *α* reduces FPR, but for *α* > 0.1, we observe little gain in FPR and a precipitous decline in recall as *α* increases. Thus, as observed earlier, *α* > 0.1 does not seem advisable. Beyond the default value, a choice of *α* = 0.01 seem desirable if more FP combinations can be tolerated. ROC curves also reveal interesting patterns in terms of the impact of ILS on the accuracy of SODA. For *α* = 0.05 and a fixed number of genes, increasing ILS increases FPR (combining species) but does not substantially impact recall. Finally, using a random resolution of the true extended species tree as the guide tree has only a small positive impact on the accuracy (Fig. S4).

### 4.2 Small simulated dataset

On the small dataset with four species and 16 populations, SODA (default) has 98% recall with both 500 and 1000 genes (Fig. 3a). Increasing the number of genes mostly reduces FPR, from 14% with 500 genes to 11% with 1000 genes. Changing *α* trades off FPR and recall in expected ways; e.g., with *α* = 0.02, recall is 100% but FPR increases to 12% for 1000 genes and 17% for 500 genes.

**Fig. 3.**
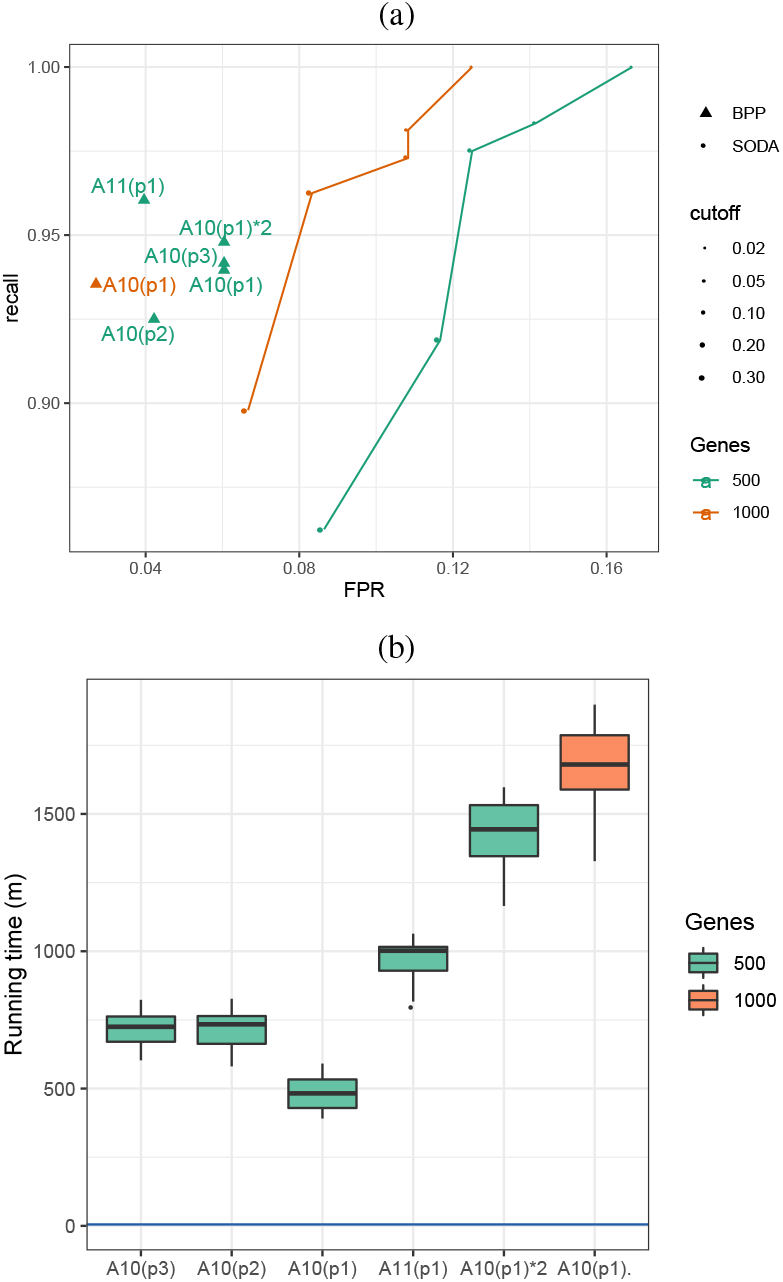
Results on the small dataset. (a) ROC curves for SODA and BPP on the small 4-taxon dataset with 500 or 1000 genes (colors) averaged over all replicates. BPP with 500 genes has several settings: three prior values for *θ_s_* are p1=IG(1.525,0.0001), p2=IG(1.525,0.001) and p3=IG(1.525,0.01). A11 indicates species tree co-estimation while A10 indicates using ASTRAL as the guide tree; A10(p1)*2 indicates doubling the number of MCMC iterations. (b) The running time of BPP with various settings. Blue horizontal line shows the running time of SODA, including gene tree estimation. Both methods are run on Intel Xeon E5-2680v3 processors; however, SODA uses one core while we ran BPP with 4 threads and 4 cores.

Compared to SODA, BPP has a lower FPR, ranging between 4% and 6% for 500 genes and 3% for 1000 genes. However, the recall of BPP is not better than default SODA and ranges between 92% and 96%, depending on the setting used. Just like SODA, increased number of genes improves FPR of BPP but not its recall. Overall, SODA-default seems to err on the side of combining individuals, while BPP tends to over-split species. Judging by the ARI (Table 1) BPP has better accuracy overall; e.g., SODA-default has an ARI of 0.78 on 500 genes while ARI of BPP ranges between 0.85 and 0.89. Overall, the parameter choices for BPP do impact accuracy but not in major ways. Doubling the number of iterations has limited to no impact and the two choices for the prior were almost identical. A third setting resulted in lower FPR but also lower recall (Fig. 3a). The only parameter that increases accuracy substantially is species tree co-estimation, which improves recall by 3.5% and reduces FPR by 0.2%.

**Table 1.** ARI on small datasets. We show mean (standard deviation) across replicates. ARI of SODA has been measured with two thresholds (0.05 and 0.1).

| Genes | SODA |  |  | BPP |  |  |  |  |
| --- | --- | --- | --- | --- | --- | --- | --- | --- |
|  | 0.01 | 0.02 | 0.05 | A10(p1) | A10(p2) | A10(p3) | A10(p1)*2 | A11(p1) |
| 500 | 0.70 (0.24) | 0.72 (0.25) | 0.75 (0.26) | 0.85 (0.20) | 0.86 (0.16) | 0.85 (0.19) | 0.86 (0.19) | 0.89 (0.15) |
| 1000 | 0.78 (0.26) | 0.79 (0.26) | 0.80 (0.23) | 0.90 (0.12) | - | - | - | - |

The slightly higher accuracy of BPP comes at a steep price in running time (Fig. 3b). BPP takes between 400 and 1900 minutes on these data, given 4 cores. In contrast, SODA never takes more than a minute. The gene tree estimation also takes a few minutes (≈ 5) for this dataset. Deviating from our default setting further increases the running time of BPP, with little impact on accuracy. For example, doubling the number of iterations results in a 3 × increase in running time and asking BPP to co-estimate the species tree results in a 2 × increase.

We next compare SODA to BPP and SpedeSTEM on the setting where for each specie, 7 individuals divided into three populations known *apriori* are given (a total of 12 populations). In this setting, FP rate of SODA is 8% on average, showing that species are sometimes combined together; in contrast, FN is 0, meaning that over-splitting does not occur (Table 2). BPP, on the other hand, outperforms SODA in terms of FP (1 to 2%), but also occasionally over-splits (FN> 0). Overall, according to ARI, BPP remains somewhat more accurate. SpedeSTEM, in comparison, has much lower accuracy; in almost all cases, it detects exactly two species (instead of four), leading to very high FP rates and low ARI. We note that in the STEM species tree inferred, the species are often non-monophyletic.

**Table 2.** ARI on small datasets with individuals assigned to populations. Three populations per species were defined randomly with 3, 2 and 2 individuals per populations. We show mean (standard deviation) across replicates. A11 indicates species tree co-estimation while A10 indicates using ASTRAL as the guide tree.

|  | SODA<br>0.05 | BPP |  | SpedeSTEM |  |
| --- | --- | --- | --- | --- | --- |
| | | A10 | A11 | $\theta = 0.1$ | $\theta = 0.01$ |
| FP | 0.08 (0.12) | 0.01 (0.02) | 0.02 (0.03) | 0.65 (0.11) | 0.78 (0.11) |
| FN | 0 | 0.01 (0.03) | 0.01 (0.02) | 0.03 (0.01) | 0.0 (0.0) |
| ARI | 0.82 (0.22) | 0.93 (0.12) | 0.89 (0.13) | 0.03 (0.11) | 0.05 (0.12) |

### 4.3 Empirical dataset

The ARI of SODA on the Protea dataset ranges from 0.61 to 0.66 with different values for *α*, given all 498 gene trees available (Table 3). This dataset includes many singleton species (12 out of 59 ingroups and 5 out of 6 outgroup species) along many non-monophyletic clades; to test the accuracy of delimitation (as opposed to species tree inference), we removed individuals that form non-monophyletic clades (19 in total) and gained better delimitation with ARI ranging from 0.72 to 0.78 on the pruned species tree.

**Table 3.** Delimitation accuracy measured using ARI for the Protea dataset with cutoff thresholds 0.01, 0.02 and 0.05. “Monophyletic species” means individuals of the species that make them non-monophyletic in the ASTRAL tree are pruned (19 in total). We run SODA on fully resolved gene trees and gene trees with branches with BS support below 5% or 10% collapsed.

| Species | Gene trees | 0.01 | 0.02 | 0.05 |
| --- | --- | --- | --- | --- |
| All species | fully resolved | 0.662 | 0.655 | 0.611 |
| | $\leq 5$ BS contracted | 0.653 | 0.654 | 0.637 |
| | $\leq 10$ BS contracted | 0.654 | 0.654 | 0.628 |
| Monophyletic species | fully resolved | 0.776 | 0.767 | 0.717 |
| | $\leq 5$ BS contracted | 0.736 | 0.775 | 0.765 |
| | $\leq 10$ BS contracted | 0.736 | 0.785 | 0.755 |

SODA-default detects 89 species and is thus over-splitting some species. Some of the split species seem to capture populations within species. For example, *Protea Acaulos* is divided into two groups that coincide with grouping based on the geographical locations of the samples (provided by the original study); these geographical separations may reflect two sub-population of this species.

We also tested collapsing low bootstrap support branches in gene trees to deal with effects of high gene tree estimation error. Collapsing very low support branches improved the results slightly with the default *α* = 0.05 (Table 3). The improvement is more clear when we prune non-monophyletic species.

#### Human

SODA results supports grouping all individuals into one species, and all the *p*-values across the tree are above 0.34. Thus, on this dataset, reassuringly, SODA avoids a false positive breakup to multiple species. While the recovery of humans (a relatively recent species) as one species may seem an easy case of species delimitation, Jackson *et al*. (2017) showed that BPP supports a four-species model with high posterior in all their 10 replicate runs, regardless of the prior used. The authors attributed this error to population structure and used this as a motivation to introduce the PHRAPL method; unlike BPP, this more advanced model did unambiguously recover humans as a single species.

#### Lizards

Statistical species delimitation using both BPP and STACEY support a speciation event at every node of the guide tree (regardless of priors chosen). SODA, similar to BPP and STACEY, detects all lineages as separate species with *p*-value≈ 0 for all branches except one branch which does not result in a false positive (Fig. S9). We note that Singhal *et al*. (2018) report that this delimitation into 13 groups does not match a clear morphological separation between species, and thus present a potential case of a cryptic species. However, it should be noted that in the light of the results from humans, a false positive delimitation by all methods cannot be ruled out.

## 5 Discussion

We designed SODA, an ultra-fast and relatively accurate method for species tree delimitation. SODA relies on frequencies of quartet topologies to decide whether each branch in a guide tree inferred from gene trees is likely to have strictly positive length, using results to infer an extended species tree, which then defines species boundaries. SODA focuses exclusively on the MSC-based species delimitation, as applicable to Eukaryotes. It is not designed for defining viral quasispecies (Domingo *et al*., 2012; Töpfer *et al*., 2014).

Our method, like many of the existing methods, is based on several strong assumptions. Most importantly, it ignores the population structure within species and does not consider gene flow. Presence of gene flow across species or population structure can lead to over-splitting for other methods like BPP, as several recent studies demonstrate (Carstens *et al*., 2013; Jackson *et al*., 2017; Sukumaran and Knowles, 2017; Leaché *et al*., 2019). Our results on the Protea dataset indicated that SODA can suffer from a similar blind-spot. We did not directly test SODA under simulation conditions with gene flow and population structure; the fact that we simulate under MSC and test under MSC can describe why our errors tend to be of over-splitting nature on simulated data. On the real Protea dataset, species were often split (often by geography), leading us to believe that SODA can be sensitive to population structure within species. However, reassuringly, SODA, unlike BPP, did not break humans into multiple species. Thus, SODA clearly has some level of robustness to population structure but is not immune from negative impacts of structure and gene flow. Note that SODA shares its shortcoming with methods purely based on MSC. Thus, we suggest that for datasets where high levels of gene flow after speciation is probable, the results of SODA should be used as a guide to enable more time-consuming delimitation using methods such PHRAPL that do consider gene flow and are more robust to structure.

In addition to gene flow and population structure, several factors need to be kept in mind when using SODA. Like other methods relying on input gene trees, the accuracy of SODA may depend on the input gene trees (Olave *et al*., 2014). It is helpful that we only rely on unrooted tree topologies, and thus, errors in branch length and rooting do not affect SODA. Plotting accuracy of SODA versus mean gene tree error across simulation conditions, except for an outlier, we didn’t detect a strong impact from gene tree error (Fig. S6). Nevertheless, errors in gene tree topologies may bias SODA towards over-splitting because gene tree error tends to increase observed discordance (Patel, 2013; Mirarab *et al*., 2014b). Moreover, the polytomy test used by SODA makes several assumptions, including the independent treatment of branches. These assumptions could in theory further impact the method in the presence of many adjacent short branches. To make sure this is not the case, our simulations used gene trees with high levels of error (Fig. S6) and included conditions with many adjacent short branches, and yet showed positive results. Nevertheless, simulations are by nature limited, and thus, further work may reveal other conditions where the method fails.

SODA also relies on a guide species tree. Luckily, given large numbers of genes, the accuracy of species trees tend to be much higher than gene trees, and Rabiee *et al*. (2019) showed that individuals of the same species often group together in ASTRAL trees. In our simulations, switching to true guide trees resulted in small improvements in accuracy (Fig. S4). Nevertheless, if the guide tree includes substantial error, SODA may suffer. For example, on the Protea dataset, several individuals were placed far from their presumed species. Assuming these individuals were correctly identified, we have to conclude the ASTRAL tree had several errors, a problem that SODA is not able to overcome. Finally, SODA requires that the analysis includes at least two individuals from each species, another factor that may limit its application to practice. Due to all these caveats, applying SODA has to be done with precautions.

We were able to compare SODA against one of the most widely-used alternatives, BPP. Previous simulation studies (e.g., Zhang *et al*., 2011; Camargo *et al*., 2012; Yang and Rannala, 2014b; Jackson *et al*., 2017) and empirical analyses (Ruane *et al*., 2013; Klein *et al*., 2016; Hotaling *et al*., 2016) have established BPP as the most accurate and preferred MSC-based delimitation method. We do not expect other Bayesian methods to be substantially more accurate than BPP (Camargo *et al*., 2012). And they are not much faster either. For example, STACEY took 7 days (Jones, 2017) on the Giarla and Esselstyn (2015) dataset with 19 individuals from 9 shrew species and 500 genes; SODA, on the same data, finished in a matter of seconds and produced identical results (Fig. S10). In the case of SpedeSTEM, it requires rooted ultrametric gene trees, which canont be inferred using the standard models of sequence evolution. Using SpedeSTEM should be combined with rooting and a rate model, which can make the analyses sensitive to errors in those steps; moreover, SpedeSTEM has been less accurate than BPP in previous analyses (Camargo *et al*., 2012). In our analyses, SpedeSTEM was the least accurate method. Perhaps the lack of accuracy is due to divergences from a strict molecular clock used in our Simphy simulations, which perhaps the SpedeSTEMv0.9 default pipeline could not overcome. STEM, used in SpedeSTEM, has been shown to have low accuracy in computing the species tree (Leaché and Rannala, 2011), especially given variations in mutational processes (Huang *et al*., 2010).

We were not able to use other methods that take gene trees as input. For example, the algorithm of O’Meara (2010) infers species delimitation either using the full gene tree likelihood calculation, which is slow Degnan and Salter (2005), or using the MDC cost Maddison (1997). However, this method (Browine) does not seem to currently have stable software support. Similarly, the method of Zhang and Cui (2010) relies on an species tree and partially labelled gene trees of individuals of several species; however, this method does not have a publicly available implementation. Older methods based on individual loci (e.g., GMYC by Pons *et al*., 2006) were not relevant to our multi-locus datasets.

The advantage of SODA over BPP, in our tests, was two-fold: much better scalability and slightly better recall. BPP cannot handle more than tens of populations while SODA can easily handle 1000 populations (used in our large simulations). However, overall, BPP was more accurate, especially when allowed to co-estimate the species tree. The relative strengths of the two methods suggest a natural way to combine them. We can first run SODA on the entire (large) dataset to obtain an initial delimitation. The results of SODA can be used to define populations and to divide the dataset into smaller subsets for a more extensive BPP analysis. This divide-and-conquer approach is what many analyses use in practice (e.g., Musher and Cracraft, 2018) using a manual curation; SODA can help automate that process.

## References

Allman, E. S., Degnan, J. H., and Rhodes, J. A. (2011). Identifying the rooted species tree from the distribution of unrooted gene trees under the coalescent. J. Math. Biol., 62, 833–862.

Camargo, A., Morando, M., Avila, L. J., and Sites, J. W. (2012). Species delimitation with abc and other coalescent-based methods: A test of accuracy with simulations and an empirical example with lizards of the liolaemus darwinii complex (Squamata: Liolaemidae). Evolution.

Carstens, B. C., Pelletier, T. A., Reid, N. M., and Satler, J. D. (2013). How to fail at species delimitation.

Coyne, J. A. and Orr, H. A. (2004). Speciation. Sinauer Associates Sunderland, MA.

Degnan, J. H. and Salter, L. A. (2005). Gene tree distributions under the coalescent process. Evolution, 59(1), 24–37.

Domingo, E., Sheldon, J., and Perales, C. (2012). Viral Quasispecies Evolution. Microbiology and Molecular Biology Reviews, 76(2), 159–216.

Ence, D. D. and Carstens, B. C. (2011). SpedeSTEM: a rapid and accurate method for species delimitation. Molecular Ecology Resources, 11(3), 473–480.

Esselstyn, J. A., Evans, B. J., Sedlock, J. L., Khan, F. A. A., and Heaney, L. R. (2012). Single-locus species delimitation: A test of the mixed yule-coalescent model, with an empirical application to Philippine round-leaf bats. Proceedings of the Royal Society B: Biological Sciences, 279(1743), 3678–3686.

Fletcher, W. and Yang, Z. (2009). INDELible: A flexible simulator of biological sequence evolution. Molecular Biology and Evolution, 26(8), 1879–1888.

Fujisawa, T. and Barraclough, T. G. (2013). Delimiting species using single-locus data and the generalized mixed yule coalescent approach: a revised method and evaluation on simulated data sets. Systematic biology, 62(5), 707–724.

Giarla, T. C. and Esselstyn, J. A. (2015). The Challenges of Resolving a Rapid, Recent Radiation: Empirical and Simulated Phylogenomics of Philippine Shrews. Systematic Biology, 64(5), 727–740.

Hebert, P. D. N., Stoeckle, M. Y., Zemlak, T. S., and Francis, C. M. (2004). Identification of Birds through DNA Barcodes. PLoS Biology, 2(10), e312.

Hotaling, S., Foley, M. E., Lawrence, N. M., Bocanegra, J., Blanco, M. B., Rasoloarison, R., Kappeler, P. M., Barrett, M. A., Yoder, A. D., and Weisrock, D. W. (2016). Species discovery and validation in a cryptic radiation of endangered primates: coalescent-based species delimitation in Madagascar’s mouse lemurs. Molecular Ecology, 25(9), 2029–2045.

Huang, H., He, Q., Kubatko, L. S., and Knowles, L. L. (2010). Sources of Error Inherent in Species-Tree Estimation: Impact of Mutational and Coalescent Effects on Accuracy and Implications for Choosing among Different Methods. Systematic Biology, 59(5), 573–583.

Hubert, L. and Arabie, P. (1985). Comparing partitions. Journal of classification, 2(1), 193–218.

Hudson, R. R. and Coyne, J. A. (2002). Mathematical consequences of the genealogical species concept. Evolution, 56(8), 1557–1565.

Huelsenbeck, J. P., Andolfatto, P., and Huelsenbeck, E. T. (2011). Structurama: Bayesian inference of population structure. Evolutionary Bioinformatics, 7, EBO–S6761.

Jackson, N. D., Carstens, B. C., Morales, A. E., and O’Meara, B. C. (2017). Species delimitation with gene flow. Systematic Biology, 66(5), 799–812.

Jones, G. (2017). Algorithmic improvements to species delimitation and phylogeny estimation under the multispecies coalescent. Journal of Mathematical Biology, 74(1-2), 447–467.

Kingman, J. F. C. (1982). On the genealogy of large populations. Journal of Applied Probability, 19(1982), 27–43.

Klein, E. R., Harris, R. B., Fisher, R. N., and Reeder, T. W. (2016). Biogeographical history and coalescent species delimitation of Pacific island skinks (Squamata: Scincidae: Emoia cyanura species group). Journal of Biogeography, 43(10), 1917–1929.

Knowles, L. L. and Carstens, B. C. (2007). Delimiting Species without Monophyletic Gene Trees. Systematic Biology, 56(6), 887–895.

Kubatko, L. S., Carstens, B. C., and Knowles, L. L. (2009). STEM: species tree estimation using maximum likelihood for gene trees under coalescence. Bioinformatics, 25(7), 971–973.

Leaché, A. D. and Rannala, B. (2011). The accuracy of species tree estimation under simulation: A comparison of methods. Systematic Biology, 60(2), 126–137.

Leaché, A. D., Fujita, M. K., Minin, V. N., and Bouckaert, R. R. (2014). Species Delimitation using Genome-Wide SNP Data. Systematic Biology, 63(4), 534–542.

Leaché, A. D., Zhu, T., Rannala, B., and Yang, Z. (2019). The Spectre of Too Many Species. Systematic Biology, 68(1), 168–181.

Maddison, W. P. (1997). Gene Trees in Species Trees. Systematic Biology, 46(3), 523–536.

Mallo, D., De Oliveira Martins, L., and Posada, D. (2016). SimPhy: Phylogenomic Simulation of Gene, Locus, and Species Trees. Systematic biology, 65(2), 334–44.

Mirarab, S. and Warnow, T. (2015). ASTRAL-II: Coalescent-based species tree estimation with many hundreds of taxa and thousands of genes. Bioinformatics, 31(12), i44–i52.

Mirarab, S., Reaz, R., Bayzid, M. S., Zimmermann, T., Swenson, M. S., and Warnow, T. (2014a). ASTRAL: genome-scale coalescent-based species tree estimation. Bioinformatics, 30(17), i541–i548.

Mirarab, S., Bayzid, M. S., Boussau, B., and Warnow, T. (2014b). Statistical binning enables an accurate coalescent-based estimation of the avian tree. Science, 346(6215), 1250463–1250463.

Mitchell, N., Lewis, P. O., Lemmon, E. M., Lemmon, A. R., and Holsinger, K. E. (2017). Anchored phylogenomics improves the resolution of evolutionary relationships in the rapid radiation of protea l. American Journal of Botany, 104(1), 102–115.

Musher, L. J. and Cracraft, J. (2018). Phylogenomics and species delimitation of a complex radiation of Neotropical suboscine birds (Pachyramphus). Molecular Phylogenetics and Evolution, 118(April 2017), 204–221.

Ogilvie, H. A., Bouckaert, R. R., and Drummond, A. J. (2017). Starbeast2 brings faster species tree inference and accurate estimates of substitution rates. Molecular biology and evolution, 34(8), 2101–2114.

Olave, M., Solà, E., and Knowles, L. L. (2014). Upstream Analyses Create Problems with DNA-Based Species Delimitation. Systematic Biology, 63(2), 263–271.

Oliveira, E. F., Gehara, M., São-Pedro, V. A., Chen, X., Myers, E. A., Burbrink, F. T., Mesquita, D. O., Garda, A. A., Colli, G. R., Rodrigues, M. T., et al. (2015). Speciation with gene flow in whiptail lizards from a neotropical xeric biome. Molecular Ecology, 24(23), 5957–5975.

O’Meara, B. C. (2010). New Heuristic Methods for Joint Species Delimitation and Species Tree Inference. Systematic Biology, 59(1), 59–73.

Pamilo, P. and Nei, M. (1988). Relationships between gene trees and species trees. Molecular biology and evolution, 5(5), 568–583.

Patel, S. (2013). Error in Phylogenetic Estimation for Bushes in the Tree of Life. Journal of Phylogenetics & Evolutionary Biology, 01(02), 110.

Pons, J., Barraclough, T. G., Gomez-Zurita, J., Cardoso, A., Duran, D. P., Hazell, S., Kamoun, S., Sumlin, W. D., and Vogler, A. P. (2006). Sequence-Based Species Delimitation for the DNA Taxonomy of Undescribed Insects. Systematic Biology, 55(4), 595–609.

Price, M. N., Dehal, P. S., and Arkin, A. P. (2010). FastTree 2 – Approximately Maximum-Likelihood Trees for Large Alignments. PLoS ONE, 5(3).

Puillandre, N., Lambert, A., Brouillet, S., and Achaz, G. (2012). Abgd, automatic barcode gap discovery for primary species delimitation. Molecular ecology, 21(8), 1864–1877.

Rabiee, M., Sayyari, E., and Mirarab, S. (2019). Multi-allele species reconstruction using ASTRAL. Molecular Phylogenetics and Evolution, 130, 286–296.

Rand, W. M. (1971). Objective criteria for the evaluation of clustering methods. Journal of the American Statistical association, 66(336), 846–850.

Rannala, B. and Yang, Z. (2003). Bayes estimation of species divergence times and ancestral population sizes using DNA sequences from multiple loci. Genetics, 164(4), 1645–1656.

Robinson, D. F. and Foulds, L. R. (1981). Comparison of phylogenetic trees. Mathematical biosciences, 53(1-2), 131–147.

Ruane, S., Bryson, R. W., Pyron, R. A., and Burbrink, F. T. (2013). Coalescent Species Delimitation in Milksnakes (Genus Lampropeltis) and Impacts on Phylogenetic Comparative Analyses. Systematic Biology, 63(2), 231–250.

Sayyari, E. and Mirarab, S. (2016). Fast Coalescent-Based Computation of Local Branch Support from Quartet Frequencies. Molecular Biology and Evolution, 33(7), 1654–1668.

Sayyari, E. and Mirarab, S. (2018). Testing for Polytomies in Phylogenetic Species Trees Using Quartet Frequencies. Genes, 9(3), 132.

Singhal, S., Hoskin, C. J., Couper, P., Potter, S., and Moritz, C. (2018). A framework for resolving cryptic species: a case study from the lizards of the australian wet tropics. Systematic Biology, 67(6), 1061–1075.

Solís-Lemus, C., Knowles, L. L., and Ané, C. (2015). Bayesian species delimitation combining multiple genes and traits in a unified framework. Evolution, 69(2), 492–507.

Sukumaran, J. and Holder, M. T. (2010). DendroPy: aPython library for phylogenetic computing. Bioinformatics, 26(12), 1569–1571.

Sukumaran, J. and Knowles, L. L. (2017). Multispecies coalescent delimits structure, not species. Proceedings of the National Academy of Sciences, 114(7), 1607–1612.

Swofford, D. L. (2001). Paup*: Phylogenetic analysis using parsimony (and other methods) 4.0. b5.

Töpfer, A., Marschall, T., Bull, R. A., Luciani, F., Schönhuth, A., and Beerenwinkel, N. (2014). Viral Quasispecies Assembly via Maximal Clique Enumeration. In Lecture Notes in Computer Science (including subseries Lecture Notes in Artificial Intelligence and Lecture Notes in Bioinformatics), pages 309–310.

Xu, B. and Yang, Z. (2016). Challenges in species tree estimation under the multispecies coalescent model. Genetics, 204(4), 1353–1368.

Yang, Z. and Rannala, B. (2010). Bayesian species delimitation using multilocus sequence data. Proceedings of the National Academy of Sciences, 107(20), 9264–9269.

Yang, Z. and Rannala, B. (2014a). Unguided species delimitation using dna sequence data from multiple loci. Molecular Biology and Evolution, 31(12), 3125–3135.

Yang, Z. and Rannala, B. (2014b). Unguided Species Delimitation Using DNA Sequence Data from Multiple Loci. Molecular Biology and Evolution, 31(12), 3125–3135.

Zhang, C., Zhang, D.-X., Zhu, T., and Yang, Z. (2011). Evaluation of a Bayesian Coalescent Method of Species Delimitation. Systematic Biology, 60(6), 747–761.

Zhang, C., Rabiee, M., Sayyari, E., and Mirarab, S. (2018). ASTRAL-III: polynomial time species tree reconstruction from partially resolved gene trees. BMC Bioinformatics, 19(S6), 153.

Zhang, J., Kapli, P., Pavlidis, P., and Stamatakis, A. (2013). A general species delimitation method with applications to phylogenetic placements. Bioinformatics, 29(22), 2869–2876.

Zhang, L. and Cui, Y. (2010). An efficient method for DNA-based species assignment via gene tree and species tree reconciliation. In International Workshop on Algorithms in Bioinformatics, pages 300–311. Springer.

