## Supplementary material for "SODA: Multi-locus species delimitation using quartet frequencies"

### 1 Supplement: proof of claims

Claim 1. *Assuming $(i)$ gene trees $\mathcal{G}$ are generated under the MSC model on an extended species tree $\mathcal{T}$, $(ii)$ the guide tree $T$ (e.g., ASTRAL tree) is a resolution of $\mathcal{T}$, and $(iii)$ the hypothesis testing has no false positive (FP) or false negative (FN) errors, the SODA algorithm returns the correct extended species tree ($\hat{\mathcal{T}}$=$\mathcal{T}$) and hence the correct delimitation under the MSC model. Additionally, it will correctly delimitate species if all species are sampled more than once.*

The proof is straightforward, and follows from the properties of the extended species tree. Under the MSC model, which is our first assumption, every branch in the guide tree that falls within a single species has zero branch length (because any resolution of the extended species tree is arbitrary). Also, internal branches of the extended species tree have non-zero length (Fig. S8). Since we assume the guide tree is a resolution of $\mathcal{T}$, the internal branches that have more than one species in their descendants should have non-zero branch lengths. If the null hypothesis of zero branch length always reject or accept the null hypothesis correctly, then, whenever we fail to reject the null hypothesis, we have a branch within a species (zero length) that needs to be contacted. And where the branch length is determined to be non-zero, the branch corresponds to a speciation event. Thus, with the assumptions of having correct gene trees, guide tree, and the hypothesis test, SODA outputs the correct extended species tree.

If each species has at least two individuals, then, for each species, there exists a bipartition such that one side of the bipartition include that species and only that species. Thus, given the correct extended species tree, and with at least two individuals per species, removing all remaining internal edges produces the correct species delimitation.

Claim 2. *Given a guide tree $T$ that resolves the tree $\mathcal{T}$, SODA incorrectly divides a species $S$ into multiple species (i.e., a false negative error) if and only if the zero length hypothesis testing results in an FP error for one of the branches under the clade defined by $S$ on $T$.*

We first prove the "if" statement. If the hypothesis test for a branch $e$ results in a false positive error, this means that $e$ had zero branch length, but the test incorrectly rejected the null hypothesis for it. In this case, the algorithm marks $e$, sister of $e$ and all ancestors of $e$ as "keep" branches and will maintain them in the extended species tree. This is equivalent to assuming that $e$ corresponds to an speciation event or descendants of $e$ form separate species from the descendants of the sister branch since this branch belongs to an internal branch of the estimated extended species tree. A false negative error happens in this case, as leaf descendants of $e$ and leaf descendants of sister of $e$ belong to the same species (since $e$ has zero branch length) and thus SODA has incorrectly divided them into two species.

Now we prove the "only if" statement. Under claim 1, we have proved that SODA returns the correct delimitation when the three conditions hold, so when guide tree $T$ is a resolution of the $\mathcal{T}$, and assuming condition 1 holds, if a species is incorrectly divided into multiple species, then there must have been at least a branch that is marked by the algorithm as "keep", but in fact should get contacted since it is a within species branch. That is only possible when the hypothesis test makes an FP error.

Claim 3. *Given a guide tree $T$ that resolves $\mathcal{T}$, SODA incorrectly combines individuals from two species $S_1$ and $S_2$ into one species (a false positive error) under one of these two conditions. 1) $S_1$ and $S_2$ each have one sampled individuals and form a cherry. 2) The hypothesis testing has an FN error for all branches of $\mathcal{T}$ below the LCA of $S_1$ and $S_2$. This condition requires FN errors for two or more branches if neither species is a singleton.*

We start with sufficiency statement. When a cherry of $S_1$ and $S_2$ each have one sampled individuals, the only branches separating $S_1$ and $S_2$ are terminal branches and we are not able to compute p-values of the null hypothesis test for terminal branches. Hence, SODA has no option but to merge the two (they won't get marked in any steps of the algorithm). For the latter case, let $e_1$ be the branch that $S_1$ individuals are pendant from and $e_2$ be the sister branch separating the subtree that $S_2$ belongs to. If the hypothesis test for a branch $e_i$ results in a false negative error, this means that $e_i$ had non-zero branch length but the test failed to reject the null hypothesis for it. In this case, the algorithm does not mark e and moves to the sister of $e_i$ (e1 or e2 in this instance). If the same condition holds for the sister branch and all branches of $T$ below the $LCA(S_1, S_2)$, then the whole subtree will not be marked and eventually the algorithm will contract those branches. This is equivalent to combining $S_1$ and $S_2$ since the only branches that will remain in the estimated extended species tree from this subtree are terminal branches forming a polytomy. A false positive error happens in this case, as $S_1$ and $S_2$ belong to the separate species and SODA has incorrectly merges them into one species.

We next prove with necessity statement. Under claim 1, we have proved that SODA returns the correct delimitation when the three conditions hold, so when guide tree $T$ is a resolution of the $\mathcal{T}$, and assuming condition 1 holds, the error could be either from a cherry of two species with one sample or an error in hypothesis test. When two or more species are merged together, this means some branches should have been marked "keep" by the algorithm but they incorrectly have been contracted. This could be the case where the test is unable to generate a p-value for the null hypothesis (terminal branches) which is only problematic when species are sampled just once, or it could be FN in the hypothesis test for the branches above those species that has resulted in the contraction of them.

### 2 Supplement: commands and parameters

simphy -rs 20 -rg 1 -rl f:1000 -sb lu:0.00001,0.00001 -sd f:0 -st f:200000 -sl f:4 -si f:10 -sp u:10000,500000 -su ln:-19,0.6931472 -hs ln:1.5,1 -hl ln:1.551533,0.6931472 -hg ln:1.4,1 -cs 9644 -o yule4 -V 3

Table S1. Parameters used in SimPhy simulation for small dataset

| Arg | Description | value |
|---|---|---|
| RS | Number of replicates | 20 |
| RL | Number of loci | 1000 |
| RG | Number of genes | 1 |
| SB | Speciation rate | 0.00001 |
| SD | Extinction rate | 0 |
| ST | Maximum tree length | 200000 |
| SL | Number of taxa | 4 |
| SI | Number of individuals per species | 10 |
| SP | Global population size | Uniform(10000,500000) |
| SU | Global substitution rate | Log normal(-19,0.6931472) |
| HS | Species specific branch rate heterogeneity rates | Log normal(1.5,1) |
| HL | Gene family specific rate heterogeneity rates | Log normal(1.551533,0.693147) |
| HG | Gene by lineage specific rate heterogeneity rates | Log normal(1.4,1) |
| CS | Random number generator seed | 9644 |

### Control File:

```
   seed = 4321
  seqfile = 16ind.sampled.phy.500g.renamed
  Imapfile = Imap.txt
  outfile = out.txt
  mcmcfile = mcmc.txt
   speciesdelimitation = 0 * fixed species tree
* speciesdelimitation = 1 0 2
   speciesdelimitation = 1 1 2 1
   speciestree = 0        * species tree NNI/SPR
   speciesmodelprior = 1
   species&tree = 16  4_0_4  4_0_7  4_0_0  4_0_8  1_0_2  1_0_3  1_0_4  1_0_8
               3_0_4  3_0_3  3_0_0  3_0_1  2_0_4  2_0_1  2_0_3  2_0_7
                        1  1  1  1  1  1  1  1  1  1  1  1  1  1  1  1
            (((4_0_4,4_0_7),(4_0_0,4_0_8)),(((1_0_2,1_0_3),(1_0_4,1_0_8))
            ,((3_0_4,(3_0_3,(3_0_0,3_0_1))),(2_0_4,(2_0_1,(2_0_3,2_0_7))))));
   diploid =   0  0  0  0  0  0  0  0  0  0  0  0  0  0  0  0
   nloci = 500  * number of data sets in seqfile
   cleandata = 0
   thetaprior = 1.525 0.0001 e  # invgamma(a, b) for theta
   tauprior = 1.525 0.0002
   locusrate = 1 5.486

   finetune =  1: 5 0.001 0.001  0.001 0.3 0.33 1.0
   print = 1 0 0 0
   burnin = 8000
   sampfreq = 2
   nsample = 200000
   scaling = 1
   threads = 4
```

**Fig. S1.** Sample control file used for running BPP with 500 genes

#### 3 Supplementary figures and tables

Table S2. Average running time of SODA (seconds) on Large dataset in different model conditions. These numbers do not include the time for inferring species tree and gene trees

| genes | 0.5M | 1M | 2M |
|---|---|---|---|
| 100 | 68.7 | 74.6 | 78.1 |
| 200 | 163.4 | 185.2 | 194.2 |
| 500 | 723.5 | 856.4 | 842.5 |
| 1000 | 1715.1 | 2071.9 | 1849.9 |

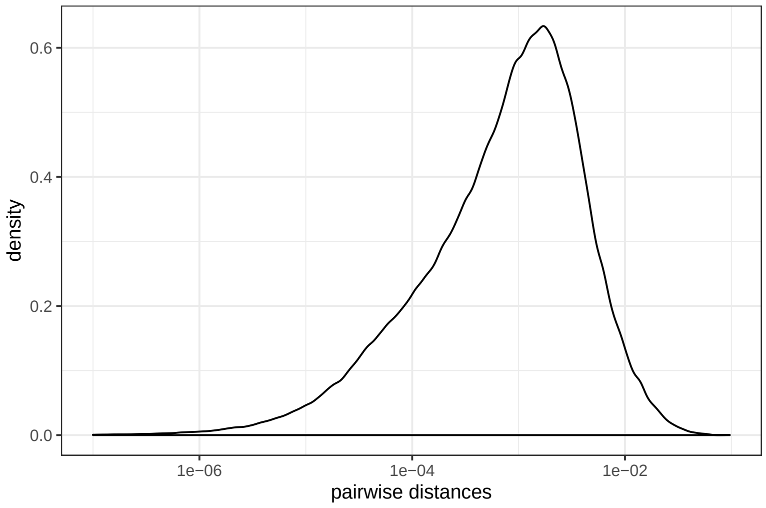


**Fig. S2.** Density plot of the pairwise distances of individuals within species in small simulated dataset computed from true gene trees in substitution units.

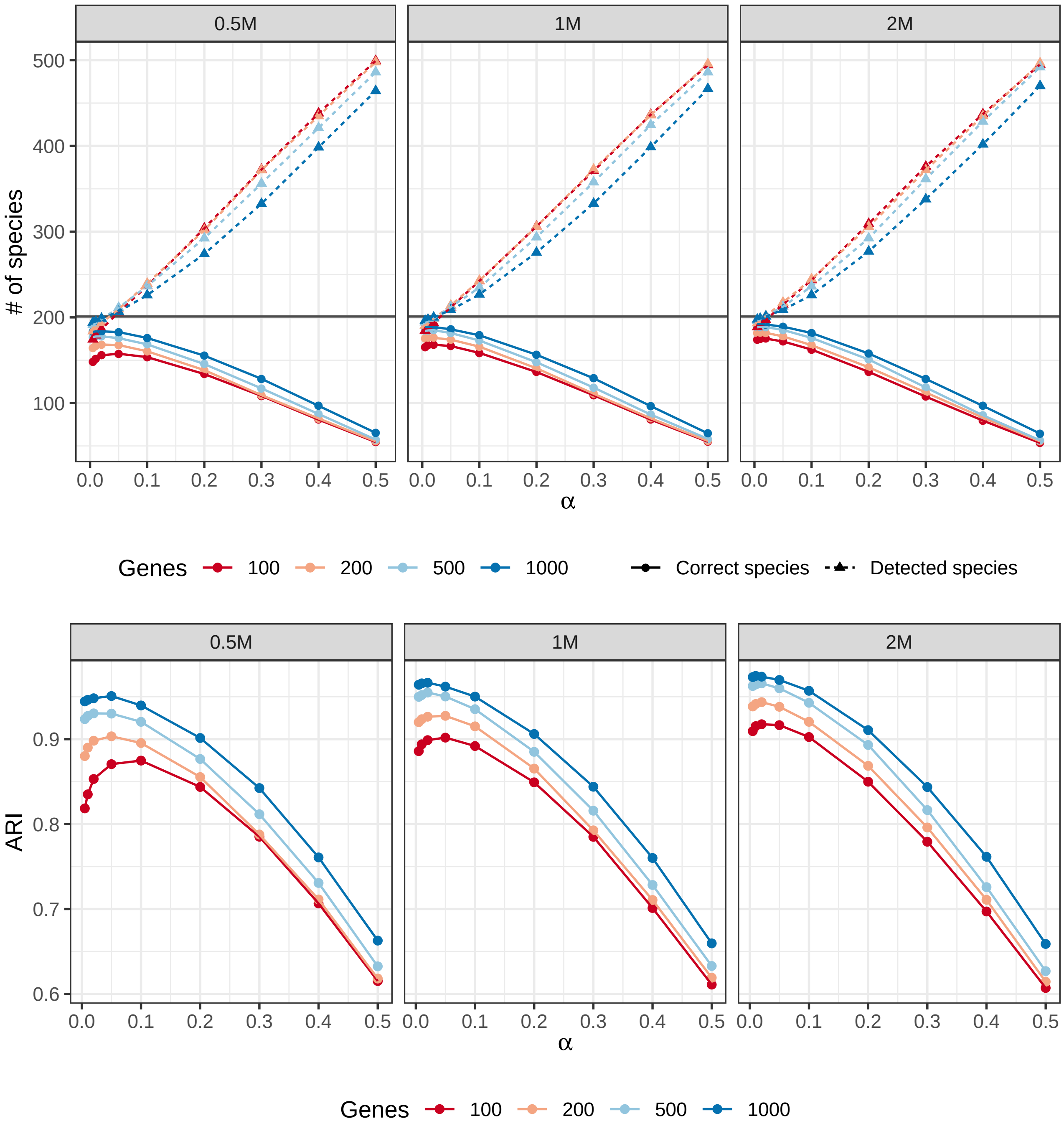


**Fig. S3. Accuracy of SODA on the large dataset.** a) We show the number of species that are completely correctly delimited (solid lines) and the total number of species found by SODA (dashed lines). Results divided into three model conditions with very high ILS (0.5M), high ILS (1M) and moderate ILS (2M) as we change $\alpha$ (x-axis) and the number of genes (colors). b) ARI (y-axis) shows the accuracy of SODA on the three model conditions with very high ILS (0.5M), high ILS (1M) and moderate ILS (2M) as we change $\alpha$ (x-axis) and the number of genes (colors).

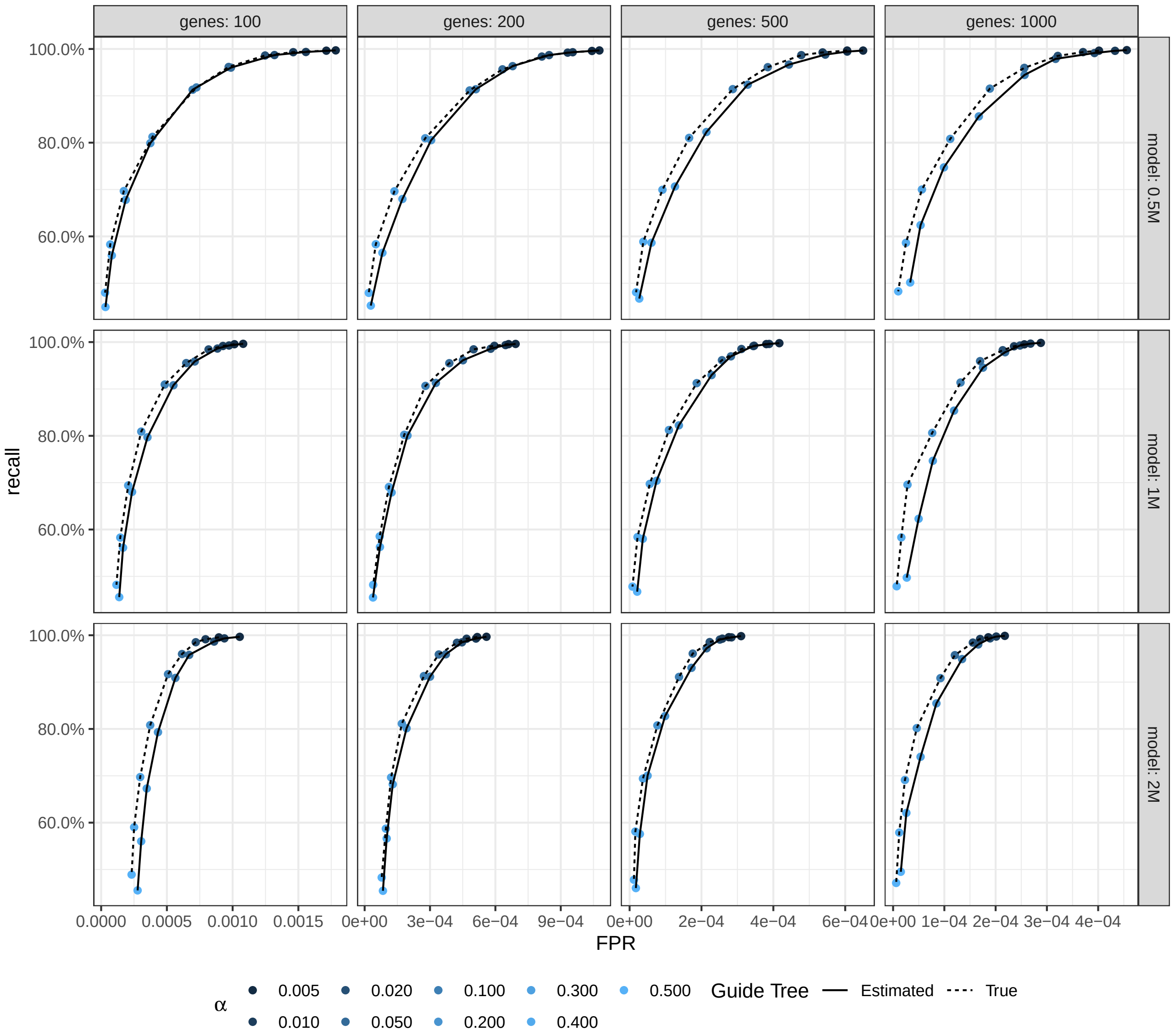


**Fig. S4. Accuracy of SODA on the large dataset.** ROC showing recall versus False Positive Rate (FPR) for all model conditions and different choices of $\alpha$ (dot size). The dotted line shows the results for true guide tree and the solid line is for estimated.

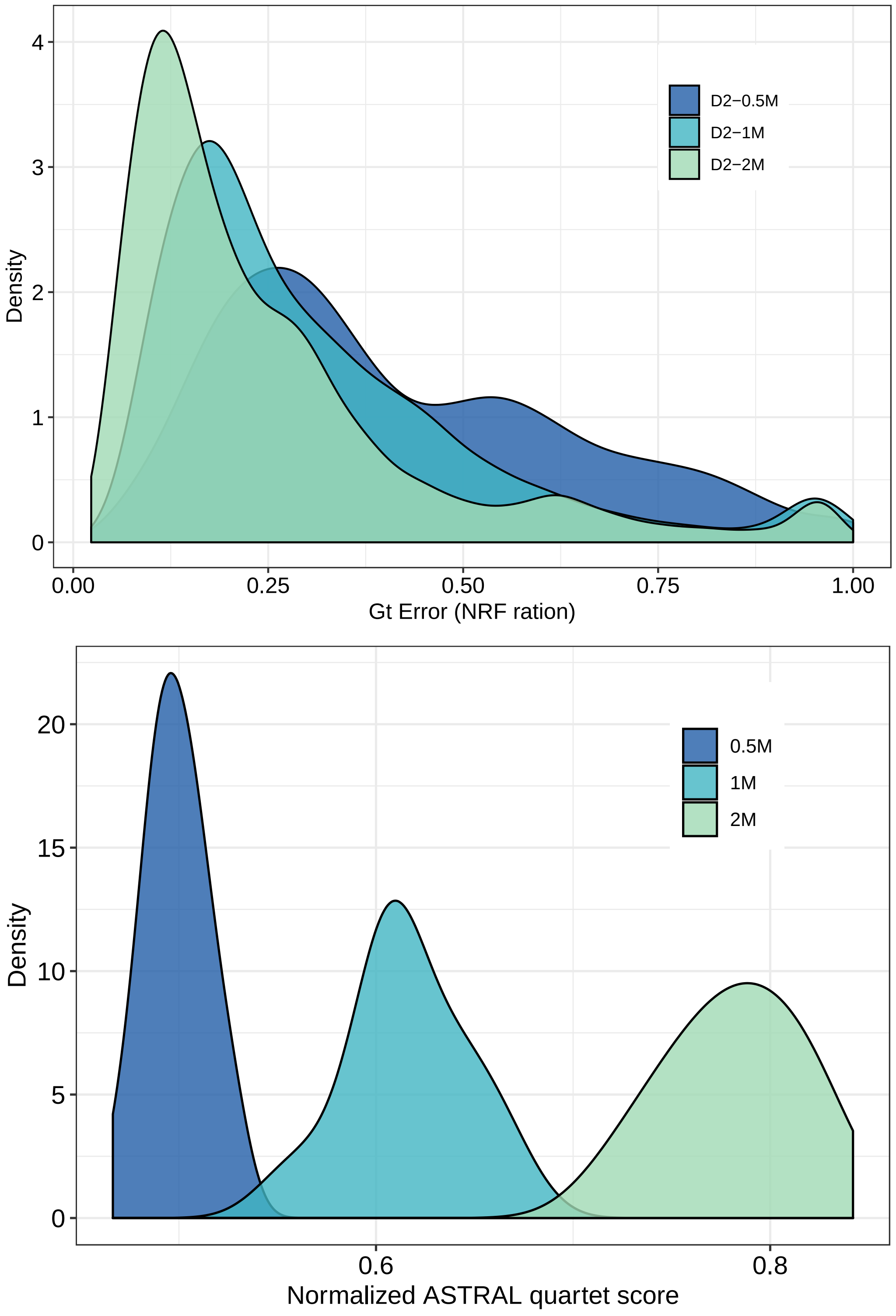


**Fig. S5.** Properties of the Large simulation dataset. We show gene tree error (top) and ILS as measured by quartet score of the species tree versus true gene trees (bottom).

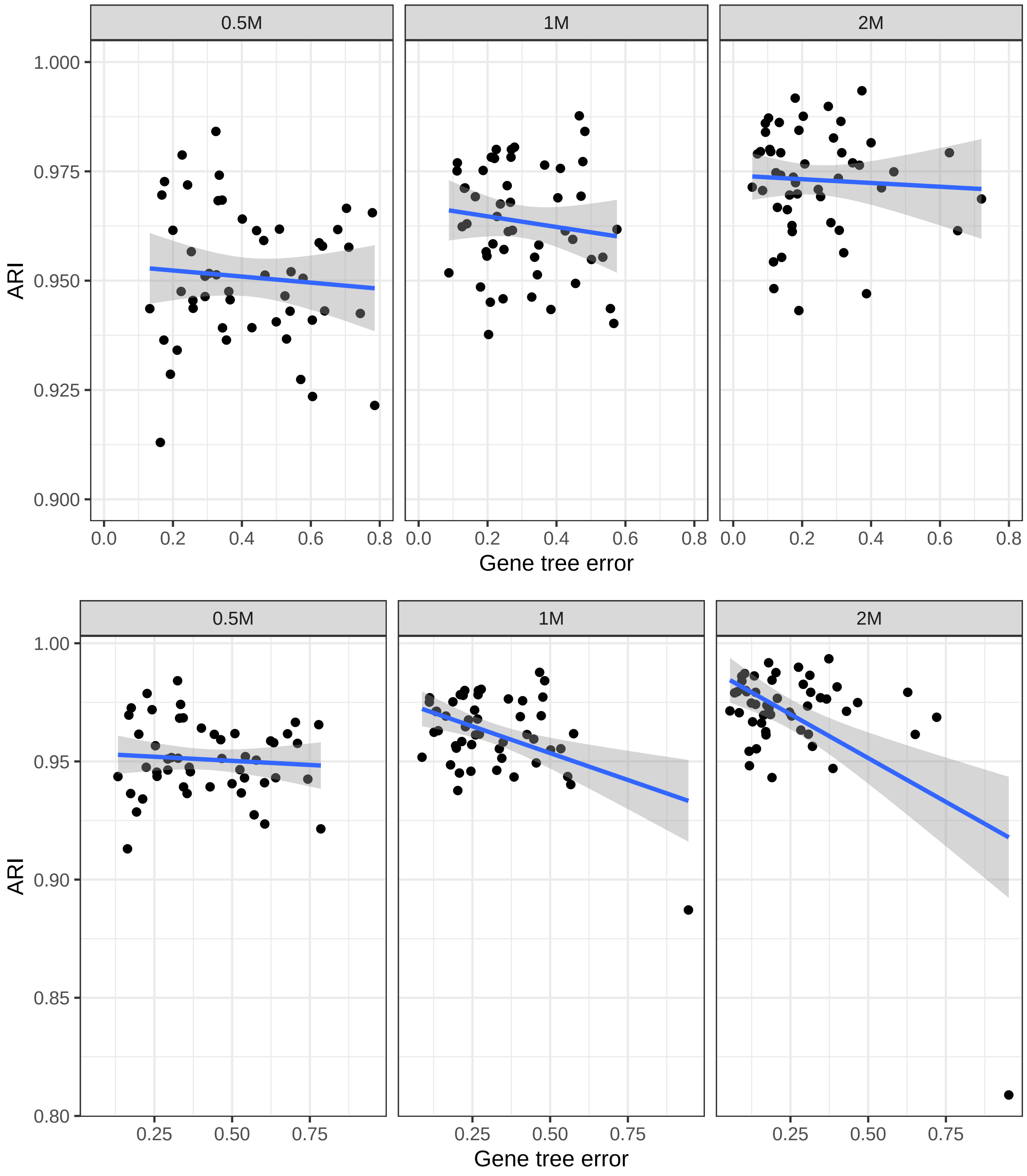


**Fig. S6.** Correlation between gene tree error and accuracy of SODA delimitation. Bottom: all replicates. Top: one outlier replicate is removed from 1M and 2M conditions.

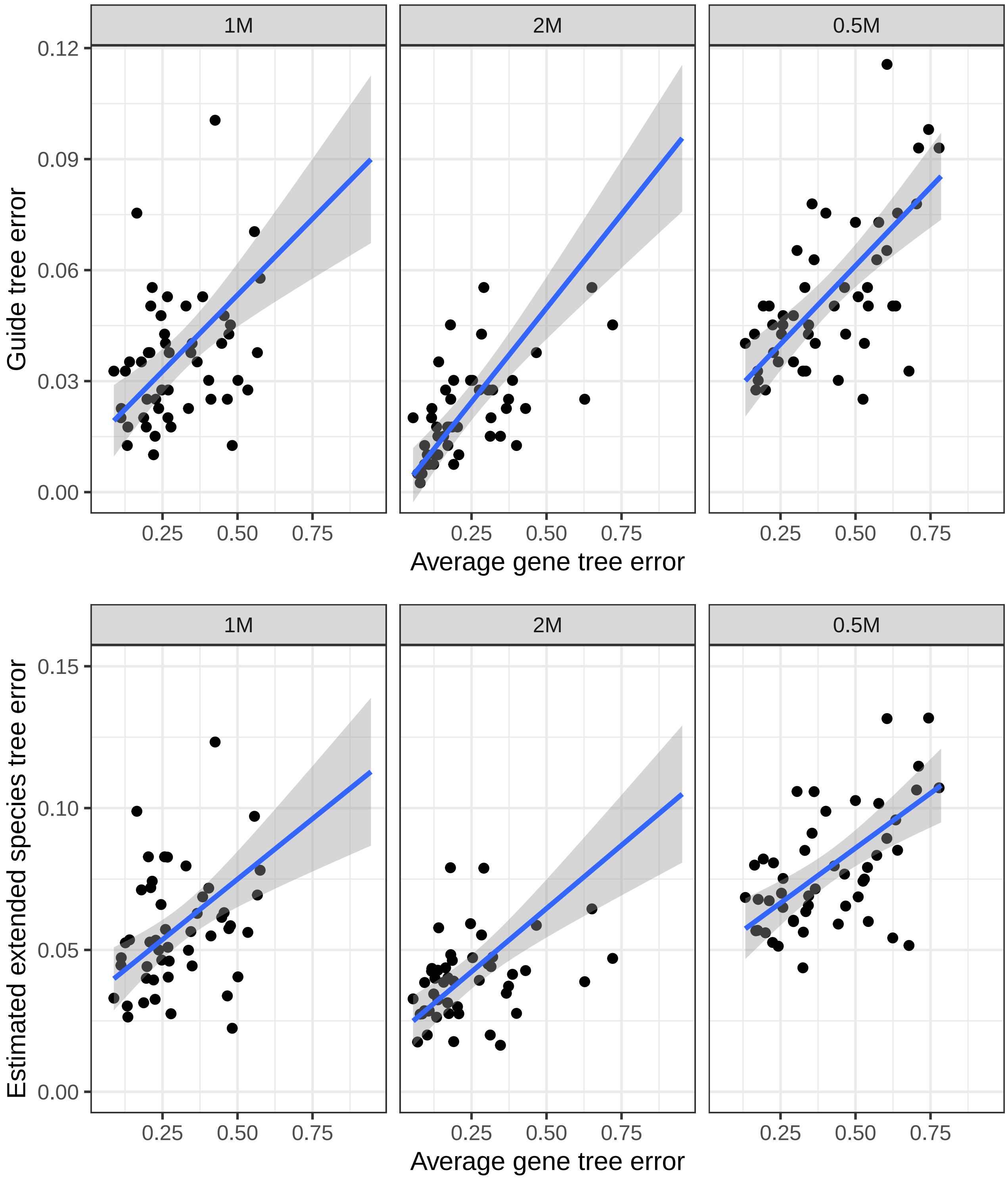


**Fig. S7.** Correlation between gene tree error and accuracy of the estimated guide tree and extended species tree. The y-axis has been limited to (0,0.12) and (0,0.15), so some outliers have been removed.

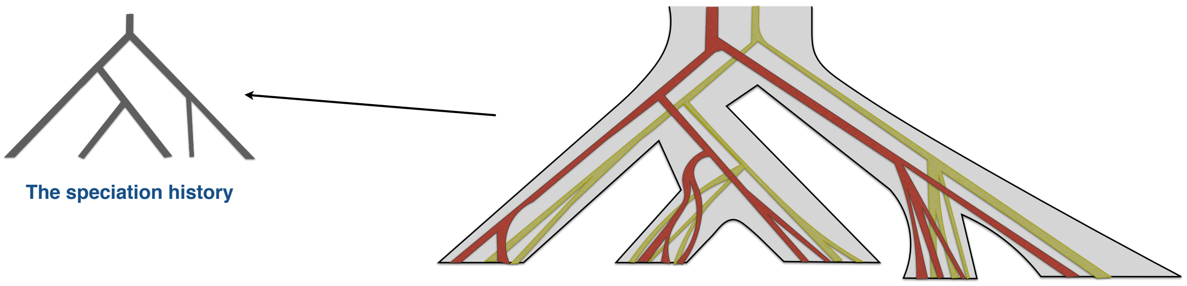


**Fig. S8.** The species tree and gene trees with individuals as tips of the tree on the right. The tree on the leftshows the true speciation history of the species that the individuals as leaves of the gene trees belong to.

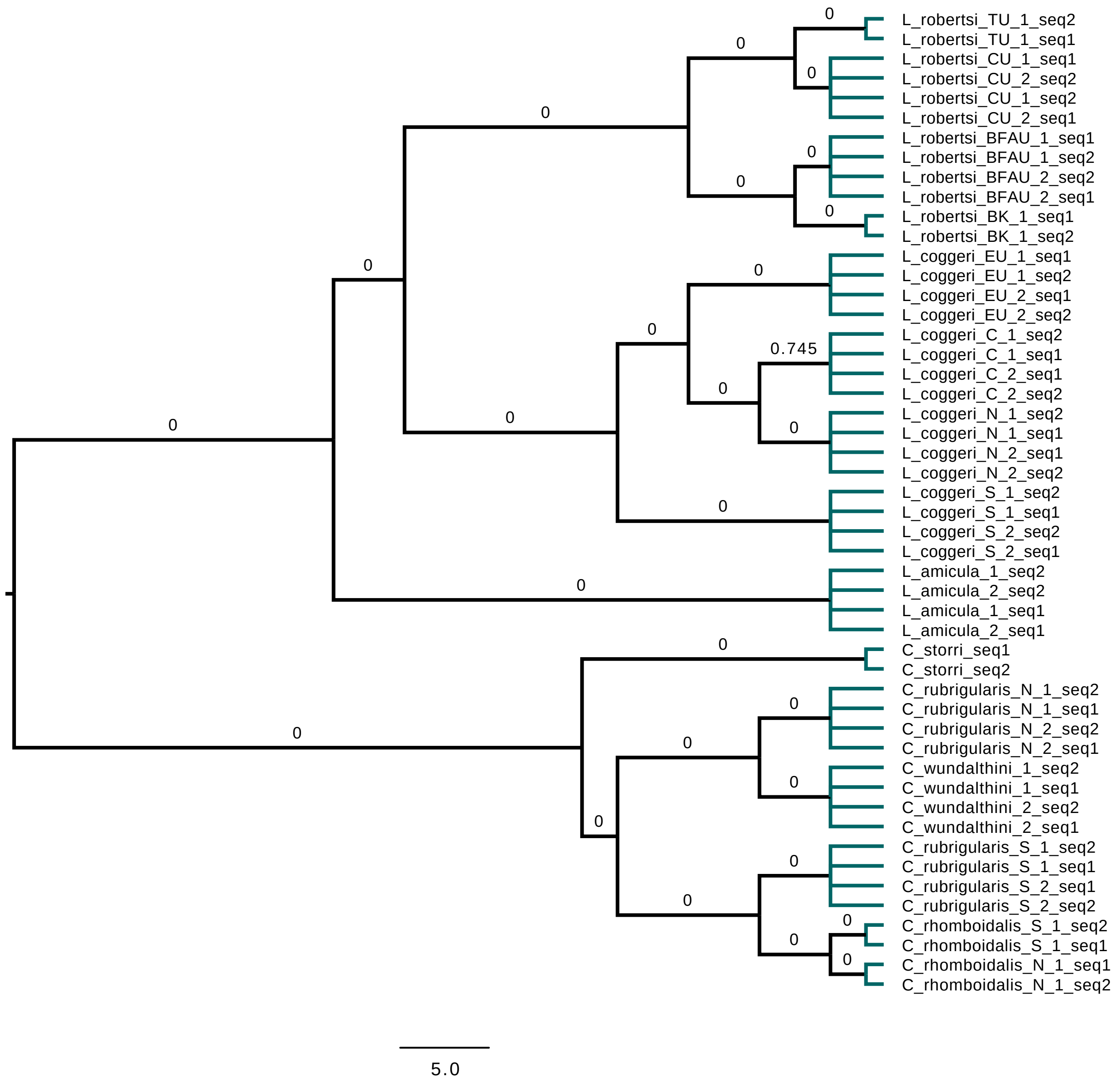


**Fig. S9.** The species tree inferred from the gene trees using multi-individual version of astral given the pre-known mapping of the individuals to species as described in Singhal et al. (2018). The values on the branches shows the p-values generated by the polytomy test.

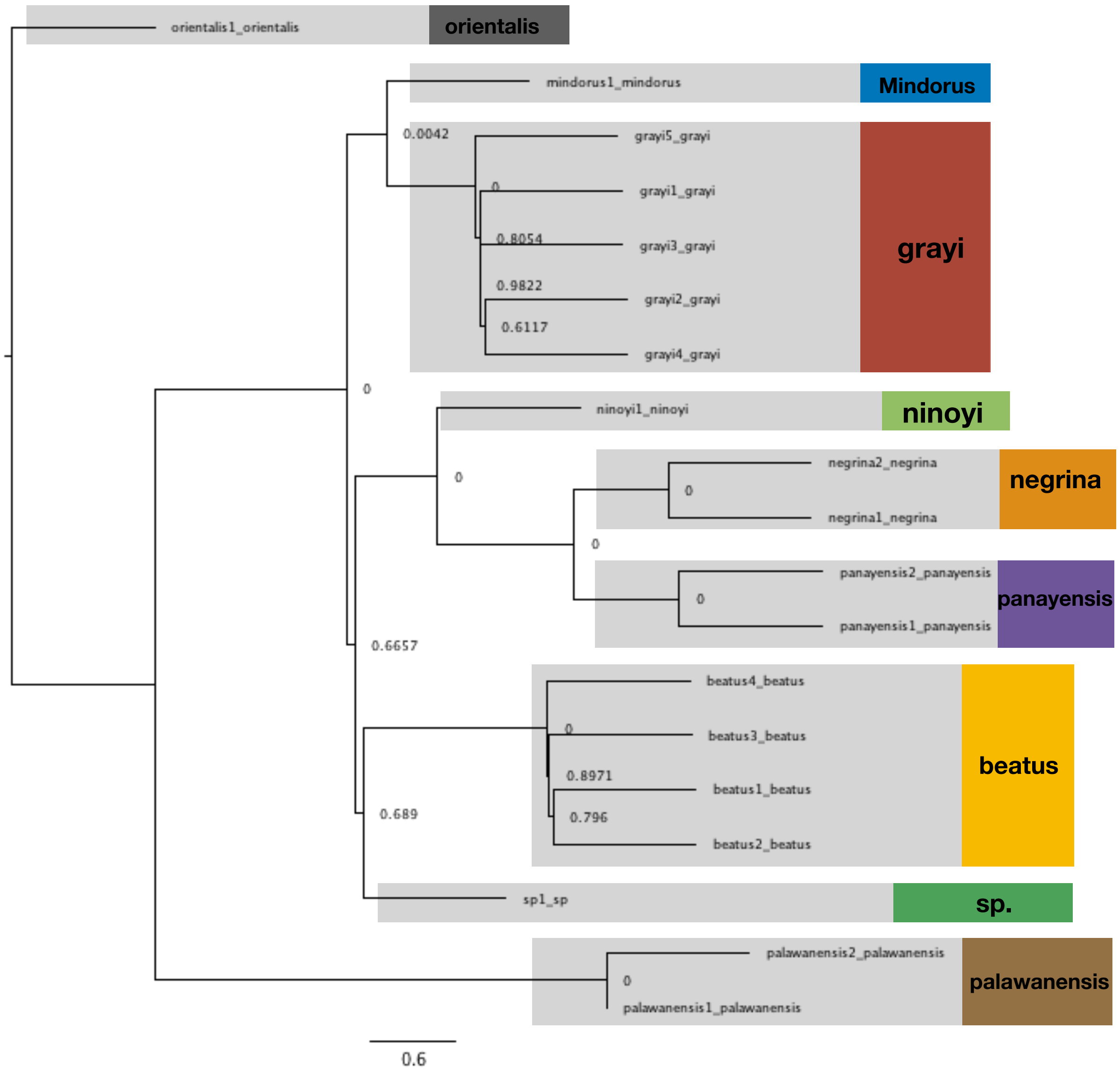


**Fig. S10.** The species tree inferred from the simulated gene trees, Sim-Matching from (Giarla and Esselstyn, 2015) , with SODA p-values put on the nodes (each number representing the p-value of the branch above that node) and the delimitation results shown as color matching the original mapping of the individuals to species.
